# Quantitative evaluation of ontology design patterns for combining pathology and anatomy ontologies

**DOI:** 10.1101/378927

**Authors:** Sarah M. Alghamdi, Beth A. Sundberg, John P. Sundberg, Paul N. Schofield, Robert Hoehndorf

## Abstract

Data are increasingly annotated with multiple ontologies to capture rich information about the features of the subject under investigation. Analysis may be performed over each ontology separately, but, recently, there has been a move to combine multiple ontologies to provide more powerful analytical possibilities. However, it is often not clear how to combine ontologies or how to assess or evaluate the potential design patterns available. Here we use a large and well-characterized dataset of anatomic pathology descriptions from a major study of aging mice. We show how different design patterns based on the MPATH and MA ontologies provide orthogonal axes of analysis, and perform differently in over-representation and semantic similarity applications. We discuss how such a data-driven approach might be used generally to generate and evaluate ontology design patterns.

## Introduction

Ontologies are widely used to characterize experimental data across all domains in the life sciences. There are now many ontologies available^1^ capturing concepts ranging from basic chemical structures^2^ through biological processes and functions^3^ to behavior^4^ and disease^5^. In biology and biomedicine, a single ontology often captures only a single kind of concept such as a process, cell type, chemical entity, phenotype, or anatomical structure, and efforts are made to ensure that the domains of description of different ontologies are orthogonal^1^.

When multiple types of data are collected in an experiment or a set of experiments, multiple ontologies may be used separately to characterize each feature. For example, annotation of zebrafish mutants with phenotypes may combine information about abnormal processes (and therefore use the Gene Ontology^3^), abnormal anatomical structures (and therefore an anatomy ontology^6^), and qualities (captured by the PATO ontology^7^): a phenotype from a zebrafish TGF β2 mutant is described as *Meckel’s cartilage chondrocyte disorganized, abnormal* being composed of the classes *Meckels Cartilage* (`ZFA:0001205`), *Chondrocyte* (`ZFA: 0009084`), *Disorganized* (`PATO:0000937`), and *Abnormal* (`PATO:00004 60`).

Ontologies are often used for data analysis^8^, for example, to facilitate enrichment analysis^9^ and semantic similarity computation^10^. When multiple ontologies are used to characterize data items, these analytical methods can be performed separately over each ontology; however, combining the different ontologies (and therefore the properties they represent) can provide improved performance for knowledge-driven data analysis approaches. However, it is often challenging to identify the right way to combine ontologies, and multiple options can exist that appear equally valid.

Ontology design patterns are adopted in constructing ontologies and describing particular phenomena within the scope of the ontology^11,12^; they are usually characterized by recurring axiom patterns, or combinations of axioms, that express certain types of knowledge and satisfy certain desiderata with regard to inferences that can be drawn from them. They may also be reused and be applied as a general strategy to structure multiple ontologies within the same or related domains. Ontology design patterns are often created around a certain structure inherent in a dataset, desired in a particular application, or considered to be intuitive by ontology users or designers^13,14^. However, in several domains, it is possible that several patterns or strategies might equally well describe the data yet permit different axes of analysis^15,16^. How to evaluate such alternative design patterns is a difficult challenge.

When evaluating ontology design patterns, we can distinguish between internal evaluation and external evaluation. An internal evaluation relies only on the ontologies and the patterns that are applied, and may involve automated reasoning to determine consistency and the number of unsatisfiable classes as well as several metrics related to the complexity of the expressed knowledge^17^. An external evaluation requires a biological hypothesis and an additional well-characterized dataset, and would involve applying the ontology design patterns to address this hypothesis; common forms of evaluation include the application of semantic similarity as a predictor for a type of biological relation.

Here, we demonstrate that we can devise and evaluate alternative design strategies using backgound knowledge from a large biological dataset, and that alternate, validated design patterns can open new axes of analysis. Specifically, we show how to combine two ontologies related to anatomy and pathology, the Mouse Anatomy Ontology (MA)^18^ and the Mouse Pathology Ontology (MPATH)^19^, through ontology design patterns. We apply several methods to evaluate the ontology design patterns through application-driven data analysis, i.e., an external evaluation of the generated ontologies.

The dataset we use for the external evaluation was derived from a very large aging study of 28 inbred strains of laboratory mice carried out at the Nathan Shock Aging Center at The Jackson Laboratory. Over their natural lifespan, mice were subject to periodic necropsy and complete histopathological workup to determine the spontaneous frequency of age-related pathological changes^20,21^.

In our analysis, first, we perform ontology enrichment analysis by strain and sex for different experimental groups, and demonstrate that different ontology design patterns yield different statistical results; we use the impact that the ontology structure has on frequently performed data analyses as additional motivation to provide quantitative measures for evaluating the design patterns. For this purpose, we compute the semantic similarities between the ontology annotations for each individual mouse and apply different clustering methods. We use a cluster purity measure with respect to our original input data and define an area under the purity curve as a quantitative measure that evaluates the quality of the different ontology design patterns. We find that there are differences between the four derived ontologies in all analysis approaches. It is well established that individual mice within an inbred (i.e., genetically homogeneous) strain exhibit a more similar spectrum of disease than mice in different strains^22,23^. We use semantic similarity measures to confirm this observation and show that some of the ontology design patterns generate significantly better results compared to using each ontology individually. Our work introduces a repertoire of quantitative ontology evaluation measures that will be useful in different applications and have the potential to improve ontology interoperability and data analysis.

## Methods

### Mouse pathology dataset

We used a dataset of spontaneous diseases of aging in the mouse^24^ available from the Mouse Phenome Database^25^. The dataset used provides 20,885 diagnoses for 1740 mice, and four different study designs. 6M involved timed necropsies at 6 months of age for a small number of strains where only selected lesions were examined. Because only a subset of data was collected, this time point is difficult to relate to the others where data collection was comprehensive, and so we do not include the 6M data in our analysis. The LONG group consisted of necropsies of moribund mice from all of the strains as they were euthanised throughout the study. The remaining groups were two cross-sectional studies at 12 and 20 months (12M and 20M) where animals from all available strains were sacrificed and necropsied together. Each diagnosis has been specified using MPATH code and MA code, other details about the diagnoses and the mice (includes scores of severity, organ, age, identification code and sex). We used 1595 mice from 28 strains; their counts are shown in table 1. Strains AKR/J, CAST/EiJ, and SJL/J were excluded because most of the animals died early in the experiment due to well-characterised severe disease or behavior. Each mouse can have multiple diagnoses and some of the mice have no diagnosis, which we will refer to as healthy mice. The experimental protocol was different in some of the groups with the 6M study only observing a subset of lesions in a preselected set of strains and the LONG group comprised of mice euthanised when moribund continuously through the study. As not all strains show high survival past 14 months, numbers necropsied for each strain are most variable at 20 months.

**Table 1.**
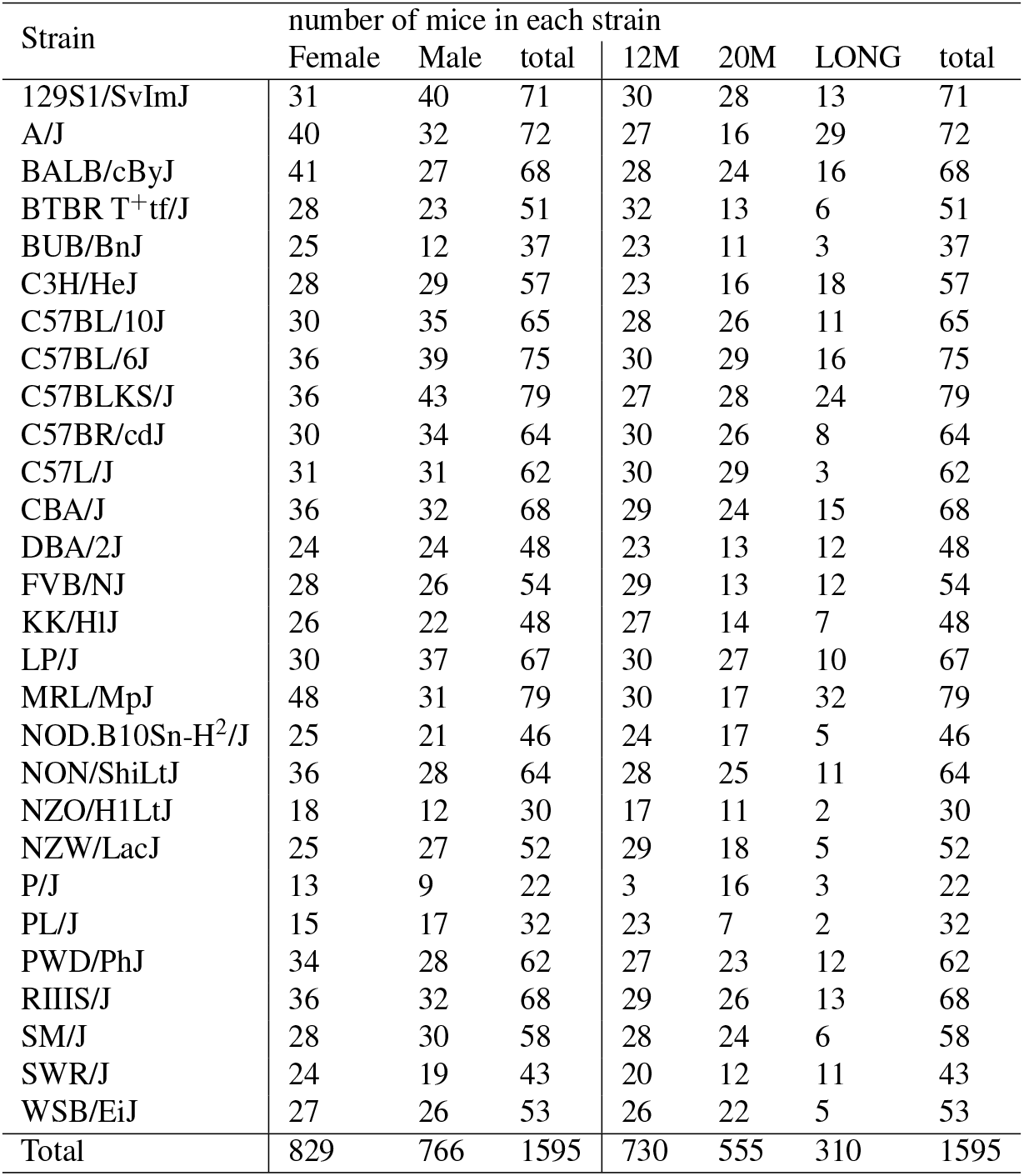
Overview of the strains used in our analysis

### Ontologies

We used two ontologies in this work, the Mouse Pathology Ontology (MPATH)^19^ and the Mouse Anatomy Ontology (MA)^18^. MPATH describes mouse pathological processes and structures. The version used was released on 2018-01-06 and contains 889 classes. MA describes adult mouse anatomy. We used the version released on 2017-02-07 and which contains 3,257 classes. As a preprocessing step in all our analyses, we added an axiom for each class in MA making them all sub-classes of a common root class, *Mouse anatomical entity*.

### Enrichment analysis

We performed enrichment analysis using the tools FUNC^26^ and OntoFunc^27^. FUNC is a software package that was developed to find significant associations between gene sets and ontological annotations in Gene Ontology, and OntoFUNC is a tool that was developed to extend the use of FUNC tool to perform enrichment analysis in ontologies other than GO.

We performed a hypergeometric test using the six different ontologies. We first applied OntoFUNC to each of the ontologies separately to generate files that will be used by FUNC. Then we generated an annotation file for each strain of mouse for each tested ontology using groovy scripts. The annotation file consists of three columns; individual mouse identifier (ID), phenotype from the ontology classes and a binary value that represents whether the mouse belongs to the strain of interest or not. Healthy mice, without phenotypes, are added by assigning them the root of the tested ontology as their phenotype. For using FUNC we specified the parameters as follows: the root for each ontology graph to be `owl:Thing`, the number of random sets to be 1,000, and ensured that each group (case and control) has at least one individual.

### Semantic similarity

We calculated mouse to mouse groupwise semantic similarity^10^ based on the existing ontologies MA, MPATH, as well as using the newly generated ontologies MAP, MAPT, PAM and PAMT. Generation of the new combined ontologies is described in Results. Briefly, MAP and MAPT are built with the MA as the primary axis of classification, and PAM and PAMT using MPATH as the primary axis of classification. MAPT and PAMT include additional axioms that base classification on the transitivity (T) of parthood relations.

We used Resnik’s similarity measure^28^ and best match average strategy implemented in the Semantic Measures Library (SML)^29^. Resnik’s similarity method is based on information content (IC) of an ontology class. The information content of a given phenotype class in the ontology is defined as the negative log of its occurrence probability^30^. As shown in Equation 1, the probability of each phenotype is calculated as the sum of each of its subclasses’ probabilities where *n_x_* is the number of occurrences of the phenotype *x* in the corpus and *x*′ ⊆ {*y*|*y* ⊑ *x*} and *N* is the is the total number of phenotype classes. The similarity between two phenotypes is then calculated as the information content of their most informative common ancestor (MICA, see Equation 3).

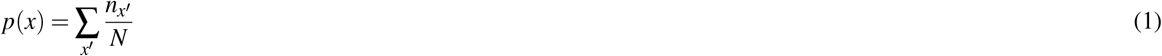

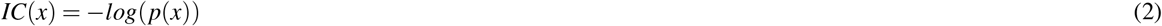

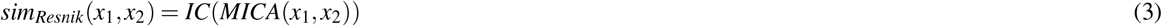

Finally, to compute mouse-to-mouse similarity, we used a best matching average method (BMA). We have two sets of ontology-based annotations, those for the first mouse and those for the second. For each annotation in either of the two annotation sets, the BMA method looks for the best match in the other set (the class with the highest similarity) and averages their similarities. In Equation 4, *m*_1_ and *m*_2_ are the number of classes associated with *mouse*_1_ and *mouse*_2_, respectively. *p*_1*i*_ and *p*_2*i*_ refer to the *i^th^* annotation of *mouse*_1_ and *mouse*_2_, respectively.

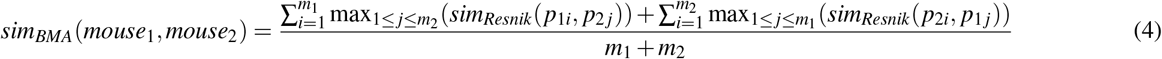

We generated mouse-to-mouse similarity scores for six groups of mice where groups are defined based on sex and age at the time of inspection. The groups are: twelve month old females (12m-F) with 372 mice, twelve month old males (12m-M) with 358 mice, twenty month old females (20m-F) with 281 mice, twenty month old males (20m-M) with 274 mice, longitudinal study females (LONG-F) with 176 mice and longitudinal study males (LONG-M) with 134 mice.

### Clustering and clustering purity

We perform clustering based on the similarity matrices generated by applying semantic similarity to each pair of mice. We applied K-medoids clustering, complete linkage agglomerative clustering, unweighted pair group method with arithmetic mean (UPGMA), and neighbor joining agglomerative clustering (NJ).

K-medoids clustering is very similar to K-means clustering but has some advantages over K-means: it does not require an observation matrix and can work directly with the similarity matrix, and it may also be less sensitive to outliers.

Hierarchical agglomerative clustering methods start by creating clusters by the number of mice. It then groups the closest clusters into one cluster one at a time and the distances between this newly generated cluster and previously existing ones are calculated. In complete linkage, the maximum distance between points in the two clusters is computed^31^. However, in UPGMA, it is calculated as the average distance between points in the two clusters^32^.

Neighbor joining^33^ is a method that is used for constructing phylogenetic trees. It starts with a star-like tree at every phase of this algorithm and tries to pick the pairs *x* and *y* with the smallest sum of branch *S_xy_* and joins them.

After clustering, we measure the quality of clusters by the cluster purity^34^. We calculate the purity based on the ground truth of mice and the strains to which they belong, by assigning each cluster to the most frequent strain in that cluster. Then, the sum of correctly assigned mice is divided by the total number of mice. Let *m*_*i*,*j*_ be the strain of mouse *j* in cluster *i*, max_i_ is the dominant strain in cluster *i*, and *M* is the total number of mice then the purity is calculated as following equations 5,6. To quantitatively compare the clustering results, we use the area under the purity curve using the trapezoidal method divided by the number of mice in the group, as shown in equation 7, where *M* is the number of mice in the group, *Purity_n_* is the purity of clusters when the number of clusters is set to *n*.

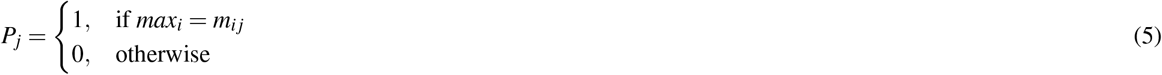

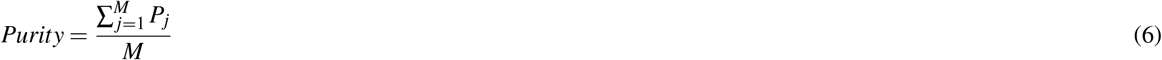

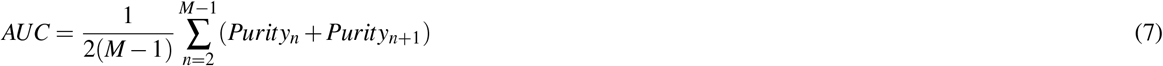

### ROC analysis

A receiver operating characteristic (ROC) curve is an evaluation measure for binary classification problems^35^. ROC curves visualize the trade-off between positives and negatives in different numerical threshold points. This allows a direct comparison between classifiers without setting a specific threshold^34^. To plot the ROC curve, we compute the true positive rate and false positive rate as shown in equations 8 and 9. This tests whether the *t* highest similar mice to a mouse would be from the same strain or not. Let *P_j_*(*t*) denote the number of mice from the top *t* most similar mice to mouse *j* that have the same strain as mouse *j* and let *M* denote the total number of mice in the tested group. We compute this for *t* = 1,2,…, *M* then plot each point that corresponds to each threshold *t* as (FPR(t),TPR(t)).

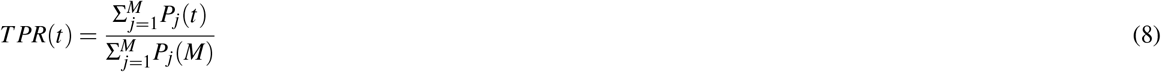

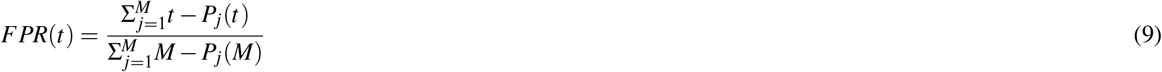

### Rank-based statistics

The ontology-based analyses that we apply in this work are enrichment and semantic similarity. The enrichment analysis provides a *p*-value for the classes that are over- and under-represented in each mouse strain, and the semantic similarity provides a similarity value for each pair of mice. We can use the *p*-values as well as the similarity score to rank classes for each strain based on their significance for over-or under-representation, and we can rank all mice for each mouse based on their pairwise similarity score. This is motivated by the practice of considering only the “most significant” or “most similar” entities as relevant, and determining how the ontology design patterns effect this kind of scenario. We can determine the strength of the effect of the different ontology design patterns based on how much the ranks (of the classes in the enrichment analysis, or the pairs of mice in the semantic similarity) change.

We apply several rank-based statistical measures, specifically Kendall’s tau correlation coefficient and the Wilcoxon rank-sum test to determine whether different ontology design patterns provide different ranks. All tests we apply are non-parametric statistical measures.

Kendall’s *τ* rank correlation coefficient is a measure of how well two sets of ranks of the same set of objects are correlated. To calculate Kendall’s *τ* we need to first calculate the number of concordant and discordant pairs. For each pair of objects this method compares the rank of those objects using the alternative ranking algorithms. If the rankings of the two objects are of the same order then they are concordant, if they are different then they are discordant. The equation to compute this is shown in 10, C is the number of concordant pairs, D is the number of discordant pairs, *t*_1_ and *t*_2_ are the number of equivalent pairs with respect to the first and second set of ranks respectively.

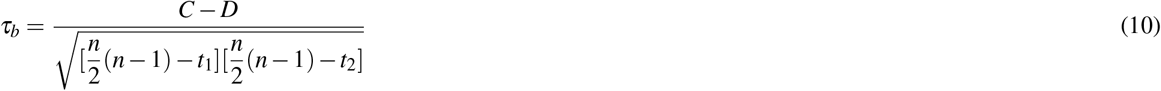

The Wilcoxon rank sum test, also known as the Mann-Whitney U-test, is a non-parametric alternative to the t-test for independent samples. We use the implementation in Matlab to perform this test.

### Implementation

We use several tools and libraries in this work. The OWL API 4.2.5^36^ is used to combine ontologies and implement our design patterns, and the FUNC 0.4.7^26^ and OntoFUNC^27^ tools are used to perform the enrichment analysis. To compute semantic similarity between mice based on their annotations, we use the semantic measure library (SML)^29^. For the quantitative and statistical analyses, we used Matlab, R and SciPy^37^ (to perform clustering, compute ROC curves and area under the ROC curves, calculate purity and rank-based statistics). All code that is required to reproduce our results, and the generated ontologies, are available at https://github.com/bio-ontology-research-group/mpath-ma.

## Results

### Ontology design patterns to combine anatomy and pathology

In the original application of MPATH, the problem of many lesions potentially occurring in many different anatomical locations was dealt with at the level of annotation, using classes from two or more ontologies to describe each lesion. This avoids the creation of all possible compound classes with the inevitable increase in class numbers, making the ontology cumbersome to use and expensive to compute over. The shortfall of this annotation-based approach is in the difficulty of using both ontologies separately in any analysis. Creating what is effectively a precomposed compound ontology obviates this problem. There are, however, different possible ways to combine the the MPATH and MA ontologies. The key problem we address is how to select the primary axis of classification, i.e., whether the classes in the combined ontology should represent anatomical entities with particular pathological lesions, or, alternatively, pathological lesions that affect particular anatomical locations. In the first case, the MA ontology will provide the natural backbone of the combined ontology’s taxonomy, while in the second case the backbone taxonomy will be provided by MPATH. A further challenge is how to incorporate information from the ontologies’ axioms in the combined ontology; for example, if a pathological lesion affects the left ventricle, we may also wish to classify this lesion as a lesion affecting the heart (and therefore utilize anatomical parthood axioms to structure the combined ontology). We combine the MPATH and MA ontologies in a data-driven way, using the OWL API^36^ to generate classes for each MPATH–MA pair observed in our mouse dataset.

To generate the first pattern, which we call MAP, we iterate through all pairs of classes from MA and MPATH observed in our mouse dataset. For each distinct pair of? *MPATH* and ?*MA* classes, we create a new class ?*MAP* defined as:

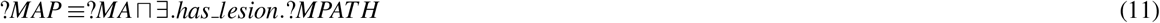

We generated the second pattern, which we call MAPT, in the same way but instead of the definition in Equation 11 we define a *MAPT* class as in Equation 12, i.e., using the parthood relation. MAP and MAPT are ontologies in which classes combine the observation that a certain anatomical entity, or its parts, have a certain lesion from MPATH. In MAPT, we reuse the `part-of` relation from the MA ontology which will then be used to infer that a lesion observed for *X* is also observed in any part of *X*. For example, if an adenoma is observed in the lungs then it is also observed in some `part-of` the respiratory system.

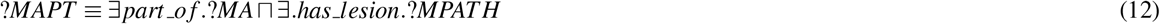

Finally, we also add a “contextualization axiom” as defined in Equation 13, which asserts that everything (that falls in the domain of our ontology) has a lesion.

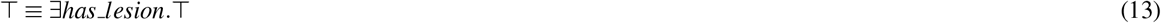

The MAP and MAPT ontologies contain 1,575 MAP classes. After generating the axioms that form the ontology, we used the HermiT reasoner^38^ to classify the ontology hierarchy.

To generate the third and fourth ontology, which we call PAM and PAMT, we define two classes from each pair of inputs to the data set in each ontology. For PAM, we define ?*PAM* class as defined in 15 and for PAMT we defined the ?*PAMT* class as defined in 14. ?*PAM* and ?*PAMT* are classes that combine the observation that a certain pathological lesion ?*MPATH* affected part of the anatomical site ?*MA*. In ?*PAMT* classes we reused the part-of relation from MA ontology for the same reason illustrated above.

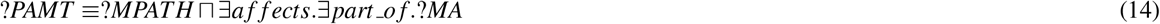

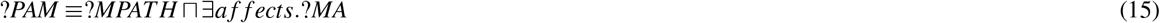

Another axiom that we added is a “contextualization axiom” axiom defined in 16, which indicates that an affect of something in any anatomical entity is a type of that thing. For example, an adenoma that affects the lung is still a type of adenoma. After PAM, PAMT and MPATH `affects` classes were asserted we inferred logical relations using the HermiT reasoner^38^. The PAM and PAMT ontologies contains 1,575 classes each.

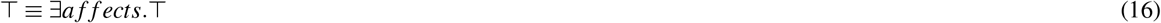

### Enrichment analysis

We performed enrichment analysis to find overrepresented lesions and anatomical sites in each strain. We used the original ontologies MA and MPATH and the newly generated ones MAP, MAPT, PAM and PAMT. Each ontology showed a characteristically different rank profile of the overrepresented classes.

We used Kendall’s rank correlation coefficient *τ* to quantify how much two ontologies, or two ontology design patterns, differ. We found that changing the primary axis between anatomy and pathology yielded highly correlated ontologies, for instance, *τ_MAP,PAM_* = 0.9990 and *τ_MAPT,PAMT_* = 0.9988. However, the correlation drops when using the transitivity of the *part_of* relation from the MA ontology, for instance, *τ_MAP,MAPT_* = 0.9130 and *τ_PAM,PAMT_* = 0.9433. Changing both the main axis of classification and using the transitivity further decreases the correlation, *τ_MAP,PAMT_* = 0.9123 and *τ_PAM,MAPT_* = 0.9145.

### Clustering purity

The driving motivation behind the generation of inbred strains of laboratory mice was that genomically identical individual mice within an inbred strain show closely related phenotypes. The phenotypic relatedness of mice within the same strain allows for the genetic analysis of genes and variants giving rise to those phenotypes^22,24,39,40^. We therefore expect that once individual mice are phenotypically annotated, these annotations should be useable in a classifier to assign each mouse to a specific strain, or at least to cluster together mice of genetically related inbred strains^23^.

Cluster purity provides a method for evaluating whether mice of the same or similar genetic background are predisposed to the same or similar lesions at same or similar anatomical sites. We first use a similarity measure to generate mouse to mouse disease similarity matrix. To eliminate confounder effects, we distinguish males and females and different time points in the study. We convert similarity matrices to distance matrices and applied four clustering methods.

The purity metric reflects how well mice of the same strain have been grouped together based solely on their phenotypic similarities, using annotation to each of the six different ontologies (i.e., MA, MPATH, MAP, MAPT, PAM, PAMT). Depending on the number of clusters, purity will naturally increase, and we determine the overall performance of the clustering task through the area under the purity curve. Table 2 shows the area under purity curves for all the group. We observe that PAM and PAMT, using the MPATH ontology as their primary axis, have the highest average AUC in the UPMGA and complete linkage methods, whereas neighbor joining and K medoids methods show MA and MAP, based on the anatomy axis do better on average in all groups.

**Table 2.** The area under cluster purity curves, based on four different clustering methods and using annotations to all six ontologies.

| Complete Linkage |  |  |  |  |  |  | UPMGA |  |  |  |  |  |  |
| --- | --- | --- | --- | --- | --- | --- | --- | --- | --- | --- | --- | --- | --- |
|  | MA | MAP | MAPT | PAM | PAMT | MPATH |  | MA | MAP | MAPT | PAM | PAMT | MPATH |
| 12m-F | 0.6324 | 0.6624 | 0.6614 | 0.6776 | 0.6740 | 0.6390 | 12m-F | 0.6093 | 0.6393 | 0.6469 | 0.6539 | 0.6619 | 0.6358 |
| 12m-M | 0.6431 | 0.6563 | 0.6572 | 0.6703 | 0.6700 | 0.6487 | 12m-M | 0.6234 | 0.6346 | 0.6423 | 0.6594 | 0.6586 | 0.6336 |
| 20m-F | 0.6532 | 0.6753 | 0.6707 | 0.6997 | 0.7036 | 0.6359 | 20m-F | 0.6215 | 0.6607 | 0.6582 | 0.6780 | 0.6699 | 0.6148 |
| 20m-M | 0.6497 | 0.6959 | 0.7144 | 0.7044 | 0.7026 | 0.6481 | 20m-M | 0.6253 | 0.6721 | 0.6903 | 0.6907 | 0.6924 | 0.6320 |
| LONG-F | 0.6654 | 0.6902 | 0.7126 | 0.6973 | 0.7049 | 0.6541 | LONG-F | 0.6496 | 0.6858 | 0.7114 | 0.6733 | 0.6832 | 0.6476 |
| LONG-M | 0.6041 | 0.6479 | 0.6329 | 0.6127 | 0.6329 | 0.5782 | LONG-M | 0.5927 | 0.6127 | 0.6209 | 0.6046 | 0.6078 | 0.5802 |
| weighted average | 0.6521 | 0.6755 | 0.6793 | **0.6863** | 0.6863 | 0.6432 | weighted average | 0.6306 | 0.6546 | 0.6641 | 0.6676 | **0.6692** | 0.6329 |
| Neighbor-Joining |  |  |  |  |  |  | K-medoids |  |  |  |  |  |  |
|  | MA | MAP | MAPT | PAM | PAMT | MPATH |  | MA | MAP | MAPT | PAM | PAMT | MPATH |
| 12m-F | 0.6618 | 0.6398 | 0.6644 | 0.6474 | 0.6249 | 0.6494 | 12m-F | 0.6380 | 0.6598 | 0.6523 | 0.6449 | 0.6468 | 0.6254 |
| 12m-M | 0.6361 | 0.6386 | 0.6312 | 0.6132 | 0.6202 | 0.6290 | 12m-M | 0.6401 | 0.6659 | 0.6596 | 0.6489 | 0.6518 | 0.6296 |
| 20m-F | 0.6817 | 0.6686 | 0.6210 | 0.6360 | 0.6434 | 0.6442 | 20m-F | 0.6366 | 0.6870 | 0.6735 | 0.6776 | 0.6834 | 0.6260 |
| 20m-M | 0.6490 | 0.6454 | 0.6397 | 0.6350 | 0.6377 | 0.6570 | 20m-M | 0.6443 | 0.6706 | 0.6838 | 0.7087 | 0.6931 | 0.6358 |
| LONG-F | 0.5719 | 0.5927 | 0.5809 | 0.5795 | 0.5832 | 0.5826 | LONG-F | 0.6780 | 0.7112 | 0.6950 | 0.7001 | 0.6938 | 0.6292 |
| LONG-M | 0.5731 | 0.5753 | 0.5641 | 0.5796 | 0.5704 | 0.5829 | LONG-M | 0.6067 | 0.6312 | 0.6325 | 0.5938 | 0.6091 | 0.5766 |
| weighted average | **0.6454** | 0.6410 | 0.6336 | 0.6287 | 0.6264 | 0.6386 | weighted average | 0.6498 | **0.6768** | 0.6708 | 0.6679 | 0.6690 | 0.6319 |

**Table 3.** Area under the ROC curve

|  | 12M-F | 12M-M | 20M-F | 20M-M | LONG-F | LONG-M | Weighted average |
| --- | --- | --- | --- | --- | --- | --- | --- |
| MA | 0.6961 | 0.7038 | 0.7412 | 0.7384 | 0.7430 | 0.7202 | 0.6962 |
| MAP | 0.7276 | 0.7287 | 0.7705 | 0.7661 | **0.7658** | **0.7362** | 0.7221 |
| MAPT | **0.7281** | 0.7340 | 0.7701 | 0.7769 | 0.7655 | 0.7345 | **0.7249** |
| PAM | 0.7149 | **0.7351** | **0.7824** | **0.7945** | 0.7465 | 0.7129 | 0.7234 |
| PAMT | 0.7126 | 0.7350 | 0.7789 | 0.7938 | 0.7494 | 0.7129 | 0.7224 |
| MPATH | 0.6950 | 0.7098 | 0.7387 | 0.7416 | 0.7044 | 0.6785 | 0.6899 |

### Semantic Similarity

Instead of using phenotype-based cluster and measuring the purity, we can also validate the hypothesis that a similar genetic background results in similar lesions directly by determining whether mice from the same stain are more phenotypically similar to each other than mice from different strains. In particular, instead of identifying how well different mouse strains separate into clusters, we are testing globally how much more similar mice in the same strain are to mice in different strains. For this purpose, we treat phenotypic similarity as a classifier that identifies mice of the same strain as positives and mice of different strains as negatives, and we can compute the true and false positive rates. A receiver operating characteristic (ROC) curve is a plot of the true positive rate as a function of the false positive rate. Figure 1 shows the ROC curves^35^ for the tested groups of mice.

**Figure 1.**
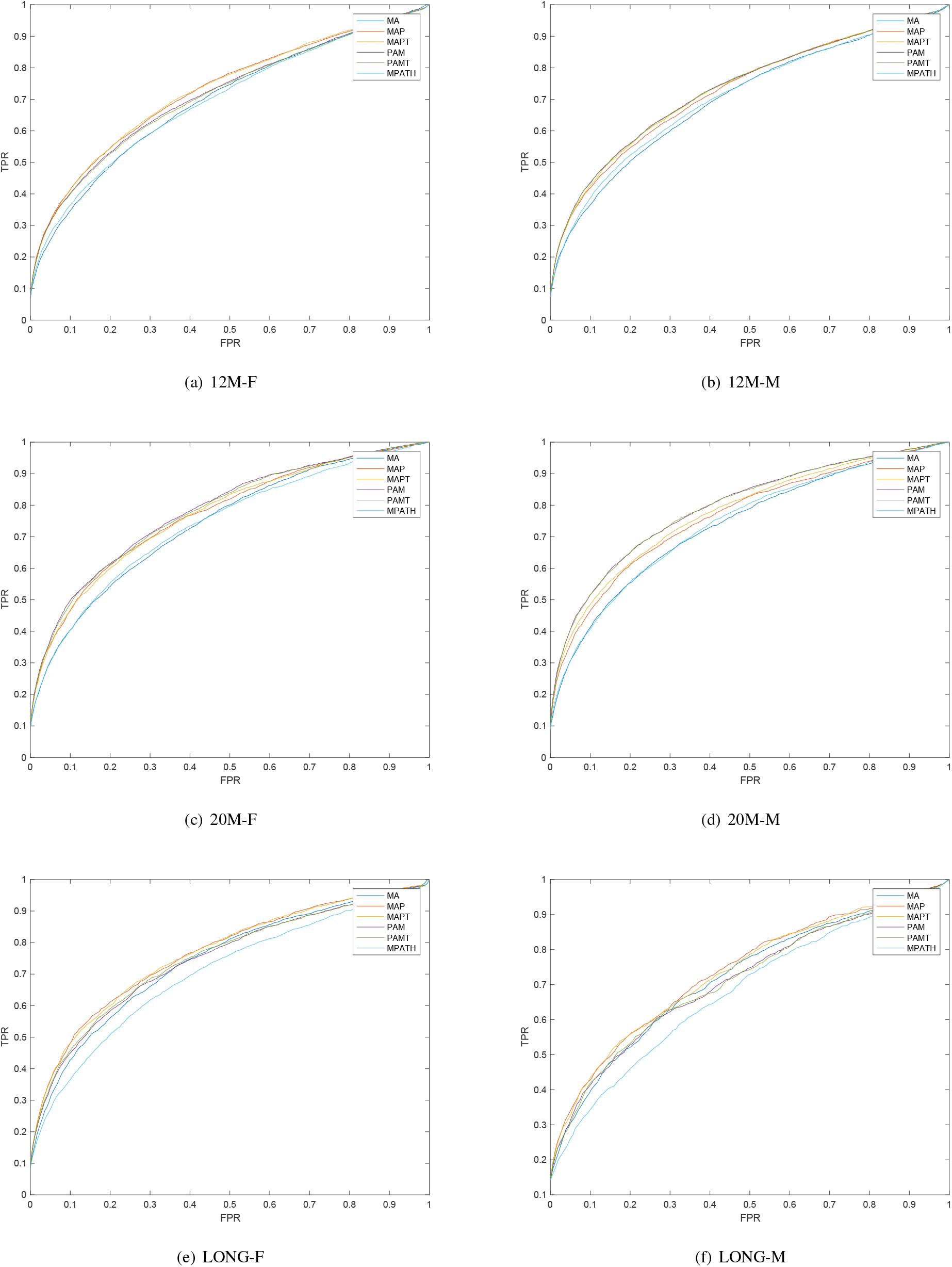
ROC curves for identifying mice from the same strain based on phenotypic similarity, and separated by the six groups of mice used in our analysis.

We further quantify the differences between the ROC curves using the area under the ROC curves (ROCAUC)^35^. We then calculated the weighted average across the four groups of mice, weighting by the number of mice in each group. We find that both MPATH and MA alone achieve a lower average AUC (0.7117 and 0.7290, respectively) than MAP (AUC 0.7454), MAPT (AUC 0.7486), PAM (AUC 0.7481), and PAMT (AUC 0.7462). The results show that the newly generated ontologies produced higher average area under the curve than MPATH and MA alone.

To test how the similarities generated by the ontologies for each pair of mice differ, we used a Wilcoxon rank sum test to test whether the differences in ROCAUC is significant. We perform the test for each pair of two ontologies and adjust p-values using Bonferroni correction. We find that there are significant differences between the new ontologies and the original ones: mice from the same strain are ranked significantly more similar than mice from different strains when using MAP, MAPT, PAM, and PAMT compared to MPATH (*p* = 4.5 · 10^−10^, *p* = 7.5 · 10^−9^, *p* = 1.1 · 10^−11^, and *p* = 9.1 · 10^−12^) as well as compared to MA (*p* = 0.02, *p* = 0.0272, *p* = 0.0159 and *p* = 0.0159, respectively). We also find that the difference between the new ontologies and MPATH is larger than the difference to MA. Among the four ontologies we generated, only the difference between MAP and MAPT as well as between MAPT and PAM/PAMT is significant (*p* = 0.021 and *p* = 0.033, respectively).

## Discussion

### Pattern-based ontology design

In the life sciences it is widely accepted that reference ontologies should cover mainly one single type of entity, and that multiple, interoperable ontologies can be used to characterize the different facets of a biological phenomenon^1^. Consequently, there is now a large set of ontologies available that can capture a wide range of phenomena^1,8^. As the ontologies are separate and often cover distinct, yet related, concepts, it is a common practice to use multiple ontologies in annotating complex datasets. For example, in annotation of protein functions, which is mainly based on the Gene Ontology (GO)^3^, additional ontologies are used to provide more complete and accurate descriptions: the Celltype Ontology (CL)^41^ or the Uberon anatomy ontology^6^ can be used to restrict certain annotations to the context of particular cell types, anatomical structures or developmental processes; the ChEBI ontology of chemical structures^2^ can provide accurate information about environmental exposures or stressors; and further ontologies can provide additional modifiers to annotations. Similarly, in the area of systems biology, it is very common to characterize models or the states of biological systems through a combination of multiple different ontologies^42^, and systematically combining these ontologies to formally describe the biological system can significantly extend the utility of individual annotations to separate ontologies^43^. Most importantly, multiple ontologies are widely combined in the area of phenotype descriptions^15^ since phenotypes can involve a wide range of morphological, environmental and processual entities, some very difficult to capture, such as lifestyle and food preferences.

Consequently, it has now become a major challenge to identify ways in which classes from multiple ontologies can be combined systematically so as to comprehensively and accurately characterize biological phenomena while maintaining the interoperability between datasets that ontologies aim to achieve. Ontology design patterns (ODPs) are an approach to provide shared, tested, and well-documented axiomatic patterns which can be applied recurrently in similar situations and therefore maintain interoperability, even when multiple ontologies are used together^11^. The application of ODPs has a wide range of purposes. The dominant ones in the biomedical sciences being patterns for standardization of content, structure and presentation, the aim being to maximize efficiency of maintenance and development^44^. Additional motivations for using ODPs are for the support of reasoning and ontology matching. Recently, motivated by the increasing importance of ontology design patterns in achieving and maintaining ontology and dataset interoperability, pattern libraries such as Dead Simple OWL Design Patterns^14^ have emerged. These libraries collect design patterns that are intended to be reused throughout the life sciences. There are often different choices in how to combine classes from different ontologies, and these choices depend on a dataset, use case, or application^45^, and some design patterns may be suited better or worse for particular applications. It has thus become a challenge to identify ways to evaluate the ontology design patterns and their utility in achieving certain outcomes; at the very least, given two choices of patterns to use (as, for example, in the area of phenotypes), it would be beneficial to determine whether the choices differ significantly or whether they likely lead to the same results.

We see one of our main contributions here as provision of a comprehensive set of evaluation methods for different ontology design patterns, and ways of comparing the effect that different ontology design patterns have on ontology-based data analysis. We use some of the most common ontology-based analysis methods in our evaluation: enrichment analysis and semantic similarity. While enrichment analysis is an exploratory method, we compare the ranks assigned through enrichment analysis to quantify how much ontology design patterns affect relative enrichment estimates. We use semantic similarity to determine how well the design patterns can reproduce a well-established biological hypothesis (i.e., that organisms with the same genotype have similar phenotypes) and quantify the effects through statistical measures, specifically, the receiver operating characteristic (ROC) curve^35^. Furthermore, we use clustering to determine if and how the different patterns make different groups of biological entities separable, and we introduce the area under the cluster purity curve as a quantitative measure.

While our aim was to provide evaluation procedures and evaluation results that are as general and generic as possible, our approach nevertheless has several crucial limitations. Most importantly, our evaluations depend on the semantic similarity measure we employ and may not generalize to other similarity measures^46^; our evaluation should be repeated if another similarity measure is chosen. Similarly, we demonstrate that the choice of the clustering algorithm changes the results, and while some general trends are observable across all algorithms we tested, the actual performance results are dependent on the algorithm. Our results do not necessarily generalize to other ontologies or even to other datasets, but demonstrate a set of methods, tools and approaches that can be employed to evaluate and test ontology design patterns.

The evaluation methods we introduce can be seen as complementary to evaluation frameworks such as OQuarE^17^ which provide quantitative statistics and measures for evaluating ontologies intrinsically. Our methods are based on an application in which ontologies are used for the analysis of a specific dataset and can therefore provide an external evaluation.

### Annotation challenges in histopathology

The formal coding of anatomic pathological observations needs to reflect the type of lesion observed together with its anatomical location and, where necessary, other characteristics such as microscopical anatomical variation, severity, or behavior. Because many lesions can occur in multiple tissues, a precomposed ontology to cover all eventualities runs into problems of combinatorial “bloat”, which makes the resulting ontology difficult to use either by humans or in computation. The solution to this challenge has been to annotate to multiple ontologies, in particular anatomy (in the case of mice to the mouse anatomy ontology MA) and pathology (from the mouse pathology ontology, MPATH), and use additional classes from PATO^47^ and other ontologies when necessary. This approach allows for the coding of almost any lesion but as the classes are used separately at the level of annotation this limits the kinds of ontology-based analysis that can be carried out. For example, even simple tasks such as counting specific lesions in given sites, e.g., determining how many mammary gland lesions of all types have been observed, becomes more challenging.

Here, we formally combine MA and MPATH into a compound ontology using two different design patterns, one in which MA is used as the ontology framework and the other MPATH. We additionally investigate the impact of introducing transitive parthood relationships into the structure of these new ontologies. Generating all possible MA and MPATH combinations is avoided by limiting the number of classes to those required to describe the dataset, plus a small number of structuring classes.

We explore design patterns for compound ontologies and evaluate alternative models of representing histopathology data based on anatomical or pathological knowledge. We are able to relate the performance of different patterns to external and independently validated concepts, namely the expectation that phenotype similarity should correlate with genotype similarity.

The first question we address is whether the compound ontologies provide a better description of the data than the single anatomy or pathology ontologies, MA and MPATH, and which of the two designs is better. The second question evaluates the impact of introducing transitivity over parthood relations into the ontology axioms. As we use completely inbred strains of mice, individuals within the same strain have an identical genotype. We utilize assessments of disease status relatedness of individual mice through semantic similarity and test globally inter- and intra-strain similarity. We find that the compound ontologies perform significantly better than the individual MA and MPATH ontologies in establishing that the mice used show closer phenotypic relatedness within, rather than between, strains. Furthermore, using clustering and evaluating cluster purity, we also find that the mice separate better in groups based on their background strain using the combined ontologies compared to using either MA or MPATH alone.

We next evaluate the primary axis of classification for the combined ontologies, being either MA or MPATH. Evaluating these different ways of combining the ontologies has significant implications not only for our dataset but also in the area of phenotype ontologies, where different phenotype ontologies have been built based on different classification axes^15,16,48,49^. In our evaluation, when we compare the performance of ranks of classes produced by the compound ontologies in enrichment analysis, we find that the primary axis of classification has little effect on the ranks (at least when using the particular aging dataset on which we rely here). We obtained similar results using the evaluation of semantic similarity measures and the clustering.

## Conclusions

Using the example of a specific large biological dataset we have shown that the data-driven generation of compound ontologies can yield powerful tools for data analysis. In addition we propose and assess comprehensive evaluation procedures for different design patterns for the resulting ontologies. We believe that these strategies for ontology generation and evaluation of ontology design patterns are generally applicable, and will be of great utility in dealing with the increasingly complex and multi-dimensional annotation of the large biomedical datasets now being widely collected.

## Acknowledgements

JPS acknowledges the support of the Ellison Medical Foundation and National Institutes of Health (AG038070-05, for the Shock Aging Center) and PNS the long term support of the Warden and Fellows of Robinson College Cambridge. RH and SMA were supported by funding from King Abdullah University of Science and Technology (KAUST) Office of Sponsored Research (OSR) under Award No. URF/1/3454-01-01 and FCC/1/1976-08-01. PNS and RH were supported by funding from King Abdullah University of Science and Technology (KAUST) Office of Sponsored Research (OSR) under Award No. FCS/1/3657-02-01.

## Author contributions statement

PNS and RH conceived of the computational experiments; SMA performed all computational experiments; SMA, PNS, RH analyzed and interpreted the results. JPS and BAS generated the aging mouse disease data. All authors reviewed and edited the manuscript.

## Additional information

### Data availability

All data and software required to reproduce our results are freely available at https://github.com/bio-ontology-research-group/mpath-ma.

## References

1. Smith, B. et al. The OBO Foundry: coordinated evolution of ontologies to support biomedical data integration. Nat Biotech 25, 1251–1255 (2007).

2. Hastings, J. et al. ChEBI in 2016: Improved services and an expanding collection of metabolites. Nucleic acids research 44, D1214–D1219 (2016).

3. Ashburner, M. et al. Gene Ontology: tool for the unification of biology. Nat. Genet. 25, 25–29 (2000).

4. Hoehndorf, R. et al. Analyzing gene expression data in mice with the Neuro Behavior Ontology. Mamm Genome 25, 32–40 (2014).

5. Kibbe, W. A. et al. Disease ontology 2015 update: an expanded and updated database of human diseases for linking biomedical knowledge through disease data. Nucleic acids research 43, D1071–D1078 (2015).

6. Mungall, C., Torniai, C., Gkoutos, G., Lewis, S. & Haendel, M. Uberon, an integrative multi-species anatomy ontology. Genome Biol. 13, R5 (2012).

7. Gkoutos, G. V., Green, E. C., Mallon, A.-M. M., Hancock, J. M. & Davidson, D. Using ontologies to describe mouse phenotypes. Genome biology 6, R5 (2005).

8. Hoehndorf, R., Schofield, P. N. & Gkoutos, G. V. The role of ontologies in biological and biomedical research: a functional perspective. Briefings Bioinforma. 16, 1069–1080 (2015).

9. Subramanian, A. et al. Gene set enrichment analysis: A knowledge-based approach for interpreting genome-wide expression profiles. Proc. Natl. Acad. Sci. United States Am. 102, 15545–15550 (2005).

10. Pesquita, C., Faria, D., Falcao, A. O., Lord, P. & Couto, F. M. Semantic similarity in biomedical ontologies. PLoS Comput. Biol 5, e1000443 (2009).

11. Gangemi, A. Ontology design patterns for semantic web content. In International Semantic Web Conference, 262–276 (2005).

12. Smith, B. et al. Relations in biomedical ontologies. Genome Biol 6, R46 (2005).

13. Mortensen, J. M., Horridge, M., Musen, M. A. & Noy, N. F. Applications of ontology design patterns in biomedical ontologies. AMIA Annu. Symp Proc 2012, 643–52 (2012).

14. Osumi-Sutherland, D., Courtot, M., Balhoff, J. P. & Mungall, C. Dead simple OWL design patterns. J. Biomed. Semant. 8, 18 (2017).

15. Gkoutos, G. V., Schofield, P. N. & Hoehndorf, R. The anatomy of phenotype ontologies: principles, properties and applications. Briefings Bioinforma. (2017). Advance access.

16. Hoehndorf, R., Oellrich, A. & Rebholz-Schuhmann, D. Interoperability between phenotype and anatomy ontologies. Bioinforma. 26, 3112–3118 (2010).

17. Duque-Ramos, A. et al. Evaluation of the OQuaRE framework for ontology quality. Expert. Syst. with Appl. 40, 2696–2703 (2013).

18. Hayamizu, T. F., Baldock, R. A. & Ringwald, M. Mouse anatomy ontologies: enhancements and tools for exploring and integrating biomedical data. Mamm Genome 26, 422–30 (2015).

19. Schofield, P. N., Sundberg, J. P., Sundberg, B. A., McKerlie, C. & Gkoutos, G. V. The mouse pathology ontology, MPATH; structure and applications. J. Biomed. Semant. 4, 1–8 (2013).

20. Yuan, R. et al. Aging in inbred strains of mice: study design and interim report on median lifespans and circulating IGF1 levels. Aging Cell 8, 277–87 (2009).

21. Sundberg, J. P. et al. Approaches to investigating complex genetic traits in a large-scale inbred mouse aging study. Vet Pathol 53, 456–67 (2016).

22. Begley, D. et al. The Laboratory Mouse, chap. Diversity of Spontaneous Neoplasms in Commonly Used Inbred Strains of Laboratory Mice, 411–426 (Academic Press, New York, NY, USA, 2012), 2 edn.

23. Beck, J. A. et al. Genealogies of mouse inbred strains. Nat. Genet. 24, 23 (2000).

24. Sundberg, J. P. et al. The mouse as a model for understanding chronic diseases of aging: the histopathologic basis of aging in inbred mice. Pathobiol. Aging & Age-related Dis. 1, 7179+ (2011).

25. Bogue, M. A. et al. Mouse phenome database: an integrative database and analysis suite for curated empirical phenotype data from laboratory mice. Nucleic Acids Res. 46, D843–D850 (2018).

26. Prüfer, K. et al. Func: a package for detecting significant associations between gene sets and ontological annotations. BMC bioinformatics 8, 41 (2007).

27. Hoehndorf, R. et al. Analyzing gene expression data in mice with the neuro behavior ontology. Mammalian genome 25, 32–40 (2014).

28. Resnik, P. Semantic similarity in a taxonomy: An Information-Based measure and its application to problems of ambiguity in natural language. J. Artif. Intell. Res. 11, 95–130 (1999).

29. Harispe, S. The semantic measures library and toolkit: fast computation of semantic similarity and relatedness using biomedical ontologies. Bioinforma. 30, 2–740 (2014).

30. Yu, G. et al. Gosemsim: an r package for measuring semantic similarity among go terms and gene products. Bioinforma. 27, 976–978 (2010).

31. Hartigan, J. A. Statistical theory in clustering. J. Classif. 2, 63–76 (1985).

32. Steinbach, M., Karypis, G. & Kumar, V. A comparison of document clustering techniques. KDD (2000).

33. Saitou, N. & Nei, M. The neighbor-joining method: a new method for reconstructing phylogenetic trees. Mol. Biol. Evol. 4, 406–425 (1987).

34. Aggarwal, C. C. Data Mining The Textbook (Springer, Yorktown Heights, New York, USA, 2015).

35. Fawcett, T. An introduction to ROC analysis. Pattern Recogn Lett 27, 861–874 (2006).

36. Horridge, M. & Bechhofer, S. The OWL API: A java API for OWL ontologies. Semantic Web 2, 11–21 (2011).

37. Jones, E., Oliphant, T., Peterson, P. et al. SciPy: Open source scientific tools for Python (2001–). URL http://www.scipy.org/. (Last accessed 27 July 2018).

38. Glimm, B., Horrocks, I., Motik, B., Stoilos, G. & Wang, Z. HermiT: An OWL 2 reasoner. J. Autom. Reason. 53, 245–269 (2014).

39. Brayton, C. F., Treuting, P. M. & Ward, J. M. Pathobiology of aging mice and gem: background strains and experimental design. Vet Pathol 49, 85–105 (2012).

40. Brayton, C. Spontaneous diseases in commonly used inbred mouse strains, vol. 3, chap. 25, 623–717 (Elsevier, Amsterdam, 2006).

41. Bard, J., Rhee, S. Y. & Ashburner, M. An ontology for cell types. Genome Biol. 6 (2005).

42. Courtot, M. et al. Controlled vocabularies and semantics in systems biology. Mol. systems biology 7 (2011).

43. Hoehndorf, R. et al. Integrating systems biology models and biomedical ontologies. BMC Syst. Biol. 5, 124+ (2011).

44. Aranguren, M. E., Antezana, E., Kuiper, M. & Stevens, R. Ontology design patterns for bio-ontologies: a case study on the cell cycle ontology. BMC Bioinforma. 9, S1 (2008).

45. Hoehndorf, R., Dumontier, M. & Gkoutos, G. V. Evaluation of research in biomedical ontologies. Briefings Bioinforma. 14, 696–712 (2013).

46. Kulmanov, M. & Hoehndorf, R. Evaluating the effect of annotation size on measures of semantic similarity. J. Biomed. Semant. 8, 7 (2017).

47. Elmore, S. et al. All in the name: A review of current standards and the evolution of histopathological nomenclature for laboratory animals. ILAR In Press (2018).

48. Köhler, S. et al. Construction and accessibility of a cross-species phenotype ontology along with gene annotations for biomedical research. F1000Research 2, 30 (2013).

49. Hoehndorf, R., Schofield, P. N. & Gkoutos, G. V. Phenomenet: a whole-phenome approach to disease gene discovery. Nucleic Acids Res 39, e119 (2011).

